## Supplementary Tables 1 to 3 and Supplementary Figures 1 to 13 for "Single-particle view of stress-promoters induction dynamics: an interplay between MAPK signaling, chromatin and transcription factors"

Victoria Wosika and Serge Pelet
Department of Fundamental Microbiology,
University of Lausanne

### Supplementary Tables

#### Supplementary Table 1. Plasmids used in this study

| Plasmid name | Description | Backbone |
|---|---|---|
| pSP264 | pSTL1 24xPP7sl | pHIS [a] |
| pSP266 | pCTT1 24xPP7sl | pHIS [a] |
| pVW200 | pGPD1 24xPP7sl | pHIS [a] |
| pVW293 | pHSP12 24xPP7sl | pHIS [a] |
| pVW294 | pGRE2 24xPP7sl | pHIS [a] |
| pVW295 | pALD3 24xPP7sl | pHIS [a] |
| pSP568 | pSTL1 24xMS2sl | pHIS [a] |
| pVW284 | pADH1 PP7ΔFG-GFPenvy tCYC1 | pSIV URA3 [b] |
| pVW300 | pTEF PP7ΔFG-GFPenvy tCYC1 | pSIV URA3 [b] |
| pVW296 | pADH1 PP7ΔFG-mCherry tCYC1 | pSIV URA3 [b] |
| pSP561 | pADH1 MS2-GFPenvy tCYC1 | pSIV URA3 [b] |
| pSP571 | pGPD Cas9 / sgRNA ($STL1_{62}$) | pRS423II [c] |
| pSP569 | pSTL1 24xPP7sl $STL1_{100-600}$ | pHIS [d] |
| pSP566 | mCherry-CaaX | pGT TRP1 [e] |

[a] Modified from Larson *et al.*[1] Integrates in the *GLT1* locus.
[b] pSIV vector from Wosika *et al.*[2].
[c] Modified from Laughery *et al.*[3]
[d] Used as repair DNA for CRISPR transformation.
[e] Modified from Wosika *et al.*[2].

**Supplementary Table 2. List of yeast strains**

All strains were constructed in the W303 background (ySP2) *MAT*a *leu2-3,112 trp1-1 can1-100 ura3-1 ade2-1 his3-11,15*. [4]

| Strain name | Relevant Genotype | Ancestor strain |
|---|---|---|
| ySP269 | HTA2-mCherry:URA3 | ySP2 |
| ySP329 | HTA2-mCherry:URA3 Hog1-GFP:HIS3 | ySP269 |
| yED215 | HTA2-mCherry:URA3 | ySP2 |
| yVW401 | HTA2-mCherry:URA3<br>pSIVu pADH1 PP7ΔFG-GFPenvy tCYC1::URA3 | yED215 |
| yVW403 | HTA2-mCherry:URA3<br>pSIVu pADH1 PP7ΔFG-GFPenvy tCYC1::URA3<br>pSTL1 24xPP7sl GLT1 tGLT1::GLT1:HIS3 | yVW401 |
| yVW428 | HTA2-mCherry:URA3<br>pSIVu pADH1 PP7ΔFG-GFPenvy tCYC1::URA3<br>pCTT1 24xPP7sl GLT1 tGLT1::GLT1:HIS3 | yVW401 |
| yVW429 | HTA2-mCherry:URA3<br>pSIVu pADH1 PP7ΔFG-GFPenvy tCYC1::URA3<br>pHSP12 24xPP7sl GLT1 tGLT1::GLT1:HIS3 | yVW401 |
| yVW430 | HTA2-mCherry:URA3<br>pSIVu pADH1 PP7ΔFG-GFPenvy tCYC1::URA3<br>pGRE2 24xPP7sl GLT1 tGLT1::GLT1:HIS3 | yVW401 |
| yVW431 | HTA2-mCherry:URA3<br>pSIVu pADH1 PP7ΔFG-GFPenvy tCYC1::URA3<br>pALD3 24xPP7sl GLT1 tGLT1::GLT1:HIS3 | yVW401 |
| yVW432 | HTA2-mCherry:URA3<br>pSIVu pADH1 PP7ΔFG-GFPenvy tCYC1::URA3<br>pGPD1 24xPP7sl GLT1 tGLT1::GLT1:HIS3 | yVW401 |
| yVW409 | HTA2-mCherry:URA3<br>pSIVu pADH1 PP7ΔFG-GFPenvy tCYC1::URA3<br>pSTL1 24xPP7sl GLT1 tGLT1::GLT1:HIS3<br>GCN5::NAT | yVW403 |
| yVW416 | HTA2-mCherry:URA3<br>pSIVu pADH1 PP7ΔFG-GFPenvy tCYC1::URA3<br>pSTL1 24xPP7sl GLT1 tGLT1::GLT1:HIS3<br>HTZ1::NAT | yVW403 |
| yVW405 | HTA2-mCherry:URA3<br>pSIVu pADH1 PP7ΔFG-GFPenvy tCYC1::URA3<br>pSTL1 24xPP7sl GLT1 tGLT1::GLT1:HIS3<br>HOT1::NAT | yVW403 |
| yVW407 | HTA2-mCherry:URA3<br>pSIVu pADH1 PP7ΔFG-GFPenvy tCYC1::URA3<br>pSTL1 24xPP7sl GLT1 tGLT1::GLT1:HIS3<br>SKO1::NAT | yVW403 |
| ySP915 | HTA2-mCherry:URA3<br>pSIVu pADH1 PP7ΔFG-GFPenvy tCYC1::URA3<br>pSTL1 24xPP7sl GLT1 tGLT1::GLT1:HIS3<br>MSN4::KAN, MSN4::NAT | yVW403 |
| yVW471 | HTA2-mCherry:URA3<br>pSIVu pADH1 PP7ΔFG-GFPenvy tCYC1::URA3<br>pGPD1 24xPP7sl GLT1 tGLT1::GLT1:HIS3<br>HOT1::NAT | yVW432 |
| yVW472 | HTA2-mCherry:URA3<br>pSIVu pADH1 PP7ΔFG-GFPenvy tCYC1::URA3<br>pGPD1 24xPP7sl GLT1 tGLT1::GLT1:HIS3<br>SKO1::NAT | yVW432 |
| ySP918 | HTA2-mCherry:URA3<br>pSIVu pADH1 PP7ΔFG-GFPenvy tCYC1::URA3<br>pGPD1 24xPP7sl GLT1 tGLT1::GLT1:HIS3<br>MSN4::KAN, MSN4::NAT | yVW432 |

| | | |
|---|---|---|
| yVW476 | HTA2-mCherry:URA3<br>pSIVu pTEF PP7ΔFG-GFPenvy tCYC1::URA3<br>pGPD1 24xPP7sl GLT1 tGLT1::GLT1:HIS3 | yVW454 |
| yVW477 | HTA2-mCherry:URA3<br>pSIVu pTEF PP7ΔFG-GFPenvy tCYC1::URA3<br>pHSP12 24xPP7sl GLT1 tGLT1::GLT1:HIS3 | yVW454 |
| yVW474 | HTA2-tdiRFP:ADE<br>pSIVu pADH1 PP7ΔFG-GFPenvy tCYC1::URA3<br>pSTL1 24xPP7sl GLT1 tGLT1::GLT1:HIS3<br>HOG1-mCherry: LEU2 | |
| ySP919 | HTA2-tdiRFP:ADE<br>pSIVu pADH1 PP7ΔFG-GFPenvy tCYC1::URA3<br>pSTL1 24xPP7sl GLT1 tGLT1::GLT1:HIS3<br>HOG1-mCherry-CaaX: TRP1 | |
| ySP921 | HTA2-tdiRFP:NAT<br>pSIVu pADH1 PP7ΔFG-GFPenvy tCYC1::URA3<br>pGPD1 24xPP7sl GLT1 tGLT1::GLT1:HIS3<br>HOG1-mCherry: LEU2 | |
| ySP922 | HTA2-tdiRFP:NAT<br>pSIVu pADH1 PP7ΔFG-GFPenvy tCYC1::URA3<br>pGPD1 24xPP7sl GLT1 tGLT1::GLT1:HIS3<br>HOG1-mCherry-CaaX: TRP1 | |
| ySP929 | HTA2-mCherry:URA3<br>pSIVu pADH1 PP7ΔFG-GFPenvy tCYC1::URA3<br>pSTL1 24xPP7sl :STL1 (clone 8) | yVW401 |
| ySP930 | HTA2-mCherry:URA3<br>pSIVu pADH1 PP7ΔFG-GFPenvy tCYC1::URA3<br>pSTL1 24xPP7sl :STL1 (clone 10) | yVW401 |
| ySP927 | MAT**a** / MATα<br>Hta2-tdiRFP:NAT / Hta2-tdiRFP :TRP<br>pSTL1 24xPP7sl GLT1 tGLT1::GLT1:HIS3 /<br>pSTL1 24xMS2sl GLT1 tGLT1::GLT1:HIS3<br>pSIVu pADH1 PP7ΔFG-GFPenvy tCYC1::URA3 /<br>pSIVu pADH1 MS2-GFPenvy tCYC1::URA3 | |
| ySP884 | HTA2-CFP:HIS3<br>HOG1-mCitrine:LEU2<br>MSN2-mCherry:URA3 | |
| ySP763 | HTA2-CFP:HIS3<br>HOG1-mCitrine:LEU2<br>pSIVu pSTL1-dPSTR-mCherry::URA3 | |
| ySP764 | HTA2-CFP:HIS3<br>HOG1-mCitrine:LEU2<br>pSIVu pHSP12-dPSTR-mCherry::URA3 | |
| yVW418 | HTA2-CFP:HIS3<br>HOG1-mCitrine:LEU2<br>pSIVu pALD3-dPSTR-mCherry::URA3 | |
| ySP766 | HTA2-CFP:HIS3<br>HOG1-mCitrine:LEU2<br>pSIVu pCTT1-dPSTR-mCherry::URA3 | |

### Supplementary Table 3. Summary of source data, strains and cell numbers

| | Strain | Stress | Replicates | Date yymmdd | Exp Num | Nb Cells |
|---|---|---|---|---|---|---|
| **Figure 1 d** | yVW403 | 0.0 M NaCl | rep1 | 181120 | 3739 | 313 |
| | yVW403 | 0.1M NaCl | rep1 | 181120 | 3745 | 404 |
| | yVW403 | 0.2M NaCl | rep2 | 181127 | 3762 | 229 |
| | yVW403 | 0.3M NaCl | rep3 | 190124 | 3849 | 201 |
| **Figure 2 a** | yVW431 | 0.2M NaCl | rep1 | 181127 | 3752 | 171 |
| | yVW428 | 0.2M NaCl | rep3 | 190118 | 3827 | 140 |
| | yVW403 | 0.2M NaCl | rep2 | 181127 | 3762 | 229 |
| | yVW430 | 0.2M NaCl | rep1 | 181127 | 3754 | 289 |
| | yVW429 | 0.2M NaCl | rep4 | 190329 | 3951 | 243 |
| | yVW432 | 0.2M NaCl | rep2 | 181127 | 3758 | 335 |
| **Figure 2 f** | yVW403 | 0.2M NaCl | rep2 | 181127 | 3762 | 229 |
| | yVW409 | 0.2M NaCl | rep3 | 190124 | 3852 | 148 |
| | yVW416 | 0.2M NaCl | rep1 | 181207 | 3796 | 175 |
| **Figure 2 g** | yVW403 | 0.2M NaCl Glucose | B | 190625 | 4076 | 248 |
| | yVW403 | 0.2M NaCl Raffinose | B | 190625 | 4076 | 275 |
| **Figure 3 a middle** | ySP329 | 0.1M NaCl | C | 190412 | 3989 | 327 |
| | ySP329 | 0.2M NaCl | C | 190412 | 3989 | 341 |
| | ySP329 | 0.3M NaCl | C | 190412 | 3989 | 311 |
| **Figure 3 a bottom** | Data from Figure 1d | | | | | |
| **Figure 3 b** | Data from Figure 2a | | | | | |
| **Figure 3 b** | Data from Figure 1d | | | | | |
| **Figure 4a** | Data from Figure 2a | | | | | |
| **Figure 4 b** | yVW430 | 0.0 M NaCl | rep1 | 181127 | 3756 | 248 |
| | yVW429 | 0.0 M NaCl | rep1 | 190222 | 3873 | 216 |
| | yVW432 | 0.0 M NaCl | rep1 | 181120 | 3743 | 214 |
| **Figure 4 c** | yVW405 | 0.2M NaCl | rep3 | 181129 | 3772 | 349 |
| | yVW407 | 0.2M NaCl | rep1 | 181120 | 3742 | 529 |
| | yVW471 | 0.2M NaCl | rep1 | 190322 | 3920 | 293 |
| | yVW472 | 0.2M NaCl | rep3 | 190405 | 3968 | 297 |
| **Figure 5** | Data from Figure 2a | | | | | |
| **Figure 6 a** | Data from Figure 1d | | | | | |
| **Figure 6 c** | yVW474 | Pulse | D | 190619 | 4056 | 168 |
| | yVW474 | Step | D | 190619 | 4054 | 159 |
| | yVW474 | Ramp | D | 190619 | 4052 | 119 |
| **Figure 7 a** | Data from Figure 2a | | | | | |
| **Sup Fig 1** | ySP884 | 0.0 M NaCl | B | 190625 | 4074 | 266 |
| | ySP884 | 0.1M NaCl | B | 190625 | 4074 | 370 |
| | ySP884 | 0.2M NaCl | B | 190625 | 4074 | 396 |
| | ySP884 | 0.3M NaCl | B | 190625 | 4074 | 291 |

| | | | | | | |
|---|---|---|---|---|---|---|
| **Sup Fig 4** | yVW403 | 0.2M NaCl | B | 200117 | 4205 | 436 |
| | ySP929 | 0.2M NaCl | A | 200117 | 4200 | 408 |
| | ySP930 | 0.2M NaCl | B | 200117 | 4203 | 483 |
| **Sup Fig 5** | yVW474 | 0.0 M NaCl | rep2 | 190503 | 4009 | 65 |
| | yVW474 | 0.1M NaCl | rep2 | 190503 | 4010 | 143 |
| | yVW474 | 0.2M NaCl | rep2 | 190503 | 4009 | 195 |
| | yVW474 | 0.3M NaCl | rep2 | 190503 | 4010 | 306 |
| **Sup Fig 6** | ySP927 | 0.2M NaCl | A | 191219 | 4191 | 89 |
| | ySP927 | 0.2M NaCl | B | 191219 | 4192 | 91 |
| | ySP927 | 0.2M NaCl | C | 191219 | 4193 | 77 |
| **Sup Fig 8** | yVW431 | 0.0 M NaCl | rep2 | 181129 | 3769 | 183 |
| | yVW428 | 0.0 M NaCl | rep2 | 181213 | 3820 | 120 |
| | yVW403 | 0.0 M NaCl | rep1 | 181120 | 3739 | 313 |
| | yVW430 | 0.0 M NaCl | rep1 | 181127 | 3756 | 248 |
| | yVW429 | 0.0 M NaCl | rep1 | 190222 | 3873 | 216 |
| | yVW432 | 0.0 M NaCl | rep1 | 181120 | 3743 | 214 |
| **Sup Fig 9 a** | Data from Figure 3a | | | | | |
| **Sup Fig 9 b** | Data from Figure 1d | | | | | |
| **Sup Fig 9 c** | Data from Figure 2a | | | | | |
| **Sup Fig 10 a** | Data from Figure 2a | | | | | |
| **Sup Fig 10 b** | Data from Figure 4b | | | | | |
| **Sup Fig 12** | yVW403 | 0.2M NaCl | rep1 | 191206 | 4164 | 269 |
| | ySP915 | 0.2M NaCl | B | 191206 | 4166 | 282 |
| | yVW432 | 0.2M NaCl | rep1 | 191206 | 4168 | 293 |
| | ySP918 | 0.2M NaCl | A | 191206 | 4169 | 310 |
| **Sup Fig 13** | yVW474 | 0.2M NaCl | C | 191217 | 4187 | 366 |
| | ySP919 | 0.2M NaCl | E | 191217 | 4188 | 257 |
| | ySP899 | 0.2M NaCl | B | 191217 | 4189 | 319 |
| | ySP922 | 0.2M NaCl | D | 191217 | 4190 | 367 |
| **Sup Fig 14** | Data from Figure 2a | | | | | |

### Supplementary Figures

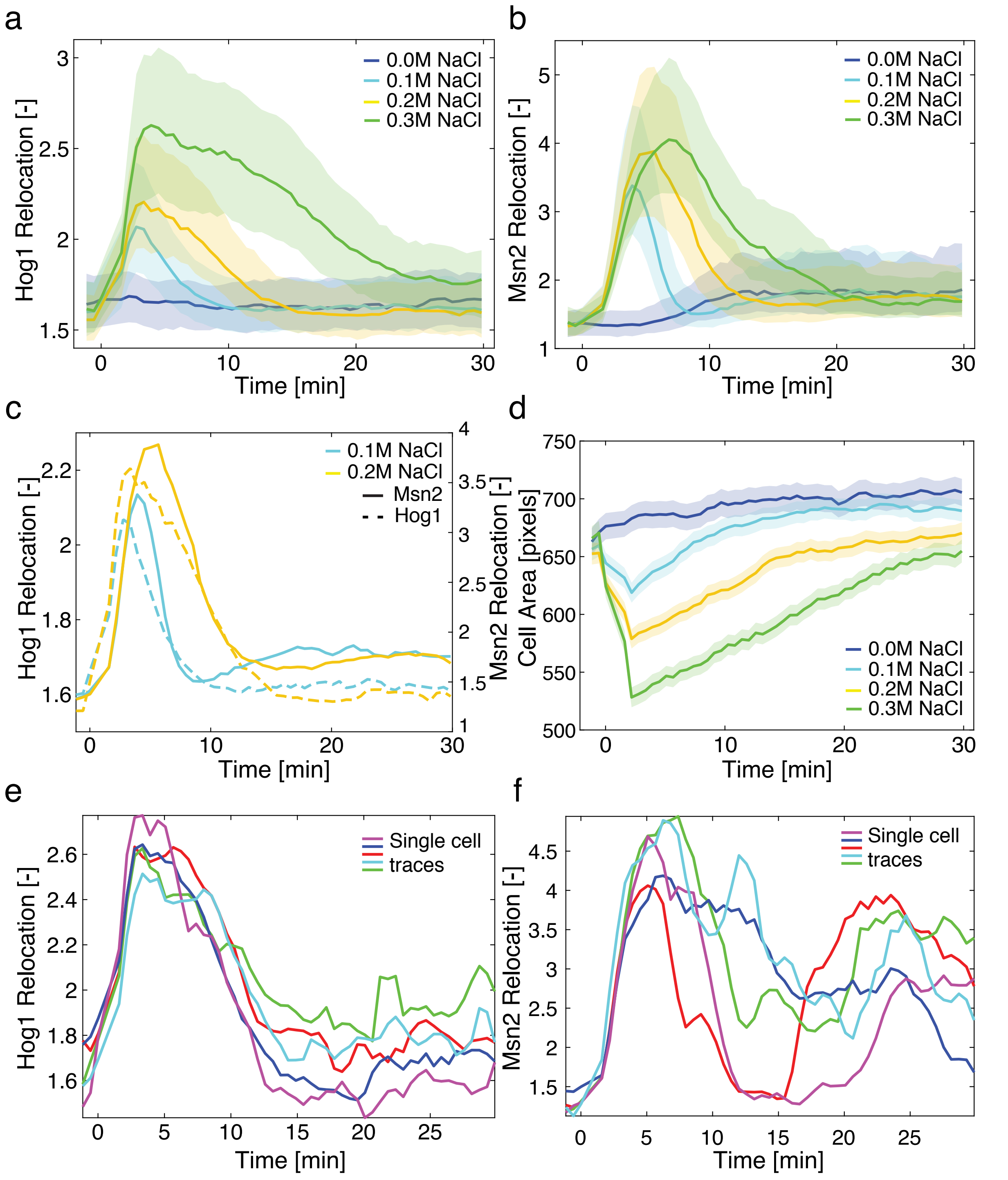


**Supplementary Figure 1. Comparison between the nuclear relocation of Msn2 and Hog1.**

**a. - c.** Strains bearing a Hta2-CFP, Hog1-mCitrine and Msn2-mCherry were stressed with various concentrations of NaCl. The nuclear relocation was quantified by the ratio in nuclear over cytoplasmic fluorescence. The median (solid line) and 25$^{th}$ -75$^{th}$ percentiles (shaded areas) are plotted for Hog1 in the yellow channel (a), Msn2 in the red channel (b) and directly compared on the same graph (c) for at least 260 cells. **d.** Change in cell size following hyper-osmotic stresses. **e. - f.** The nuclear relocation traces from the same single cells for Hog1 (e) and Msn2 (f) are plotted. These cells were selected in the population because they display a strong re-entry of Msn2 in the nucleus following the first pulse of activity. This second phase in the response is absent from the Hog1 dynamics.

a

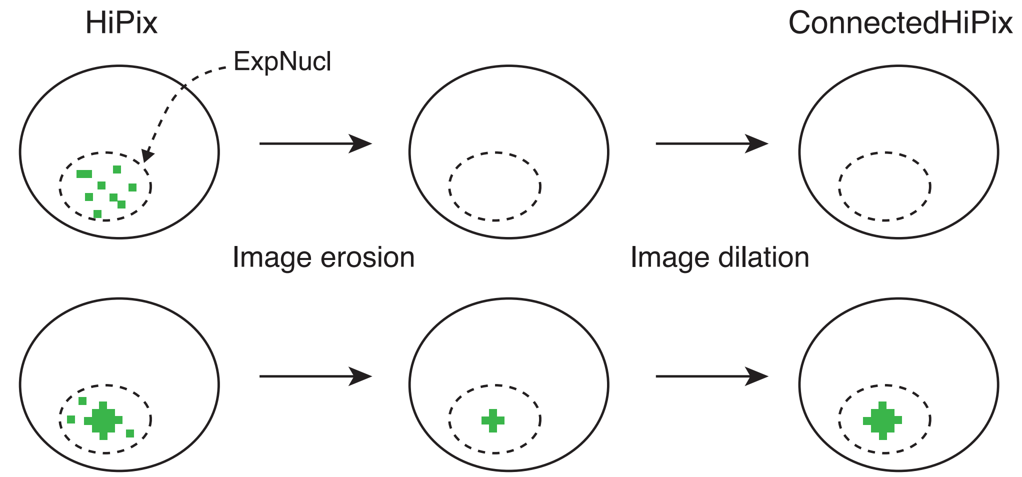


b

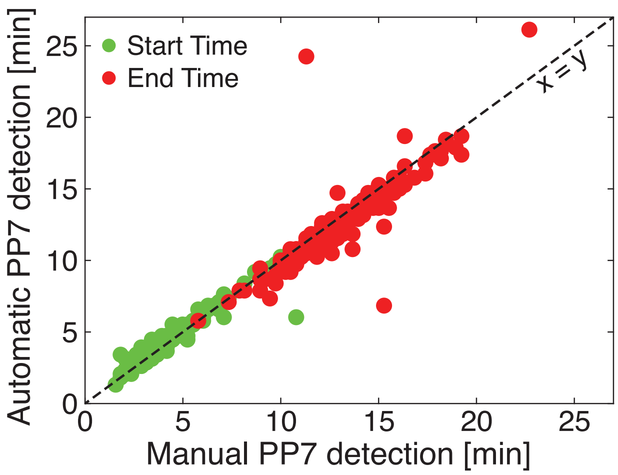


c

| | | Manual Curation | |
|---|---|---|---|
| | | **PP7 Positive** | **PP7 Negative** |
| **Automated measurement** | **PP7 Positive** | 193 | 1 |
| | **PP7 Negative** | 1 | 43 |

**Supplementary Figure 2. Quantification of the PP7 traces with the ConnectedHiPix feature.**
**a.** Scheme describing the process used to generate the ConnectedHiPix feature from the 20 highest intensity pixels (HiPix) in the ExpNucl object (Nucleus expanded by 5 pixels). **b.** Comparison between a manual curation of Start Times (green) and End Times (red) by visual inspection of the cells and automated quantification by the ConnectedHiPix measurement. **c.** Table displaying the number of PP7 positive and negative cells obtained by manual versus automated quantification.

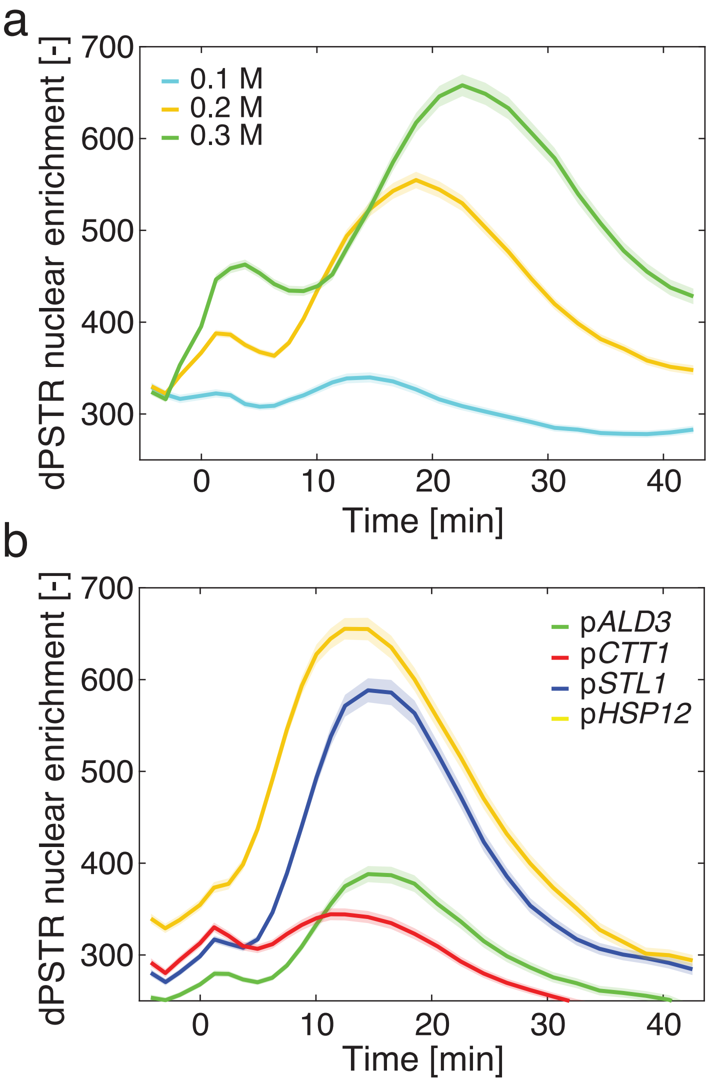


**Supplementary Figure 3. Dynamic protein synthesis translocation reporter (dPSTR) induction upon osmotic stress.**
The dPSTR reporter allows to by-pass the slow maturation time of protein expression reporters. A constitutively expressed FP is functionalized by a leucine zipper. The compatible zipper is under the control of the inducible promoter of interest and coupled to two strong Nuclear Localization Signals (NLS) motifs. When the pair of leucine zippers interact, the enrichment of the FP in the nucleus allows to quantify the level of induction of the promoter[5]. **a.** p*STL1*-dPSTR$^{R}$ nuclear enrichment (nuclear fluorescence minus cytoplasmic fluorescence) for three different stress levels. **b.** dPSTR$^{R}$ nuclear enrichment for 4 different promoters following a 0.2M NaCl stress. In all graphs, the solid line represents the mean difference and the shaded area represents the s.e.m.

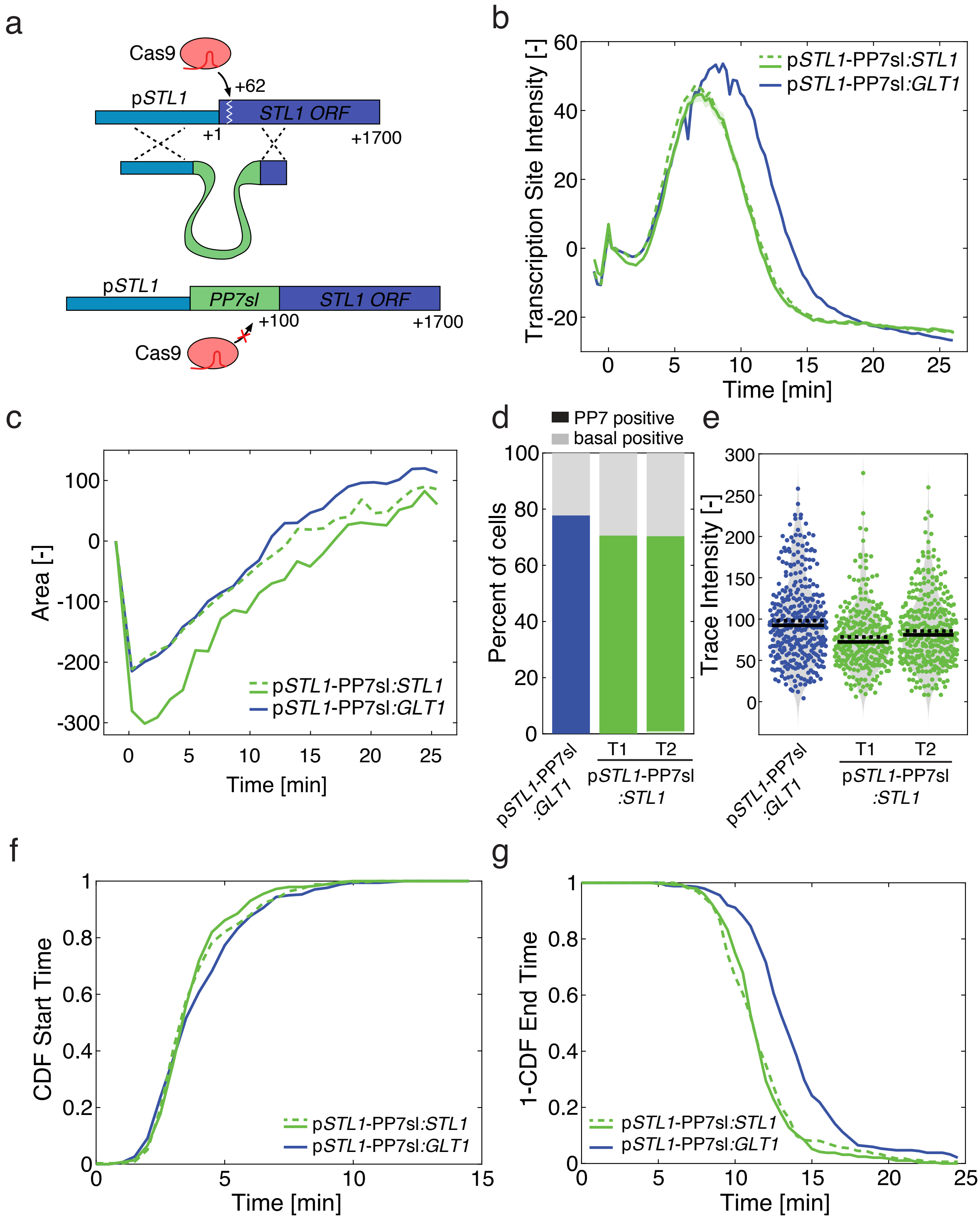


**Supplementary Figure 4. Monitoring transcription from the endogenous *STL1* locus** **a.** Scheme describing the integration of the 24xPP7sl at the *STL1* locus by CRISPR-Cas9. The sgRNA recognizes the PAM motif 62bp upstream of the start codon. The DNA break is repaired by homologous recombination from a DNA fragment that contains homology with the *STL1* promoter and 500 bp from the *STL1* ORF starting at position 100. **b.** Transcription site intensity arising from the p*STL1* promoter upon 0.2M NaCl stress. **c.** Cell size adaptation dynamics following hyper-osmotic for the experiment presented in b. **d.** Percent of PP7 positive cells. **e.** Maximum intensity of single-cell traces. **f.** and **g.** Dynamics of transcription onset and shut off represented with the CDF of Start Times (f) and 1-CDF of End Times (g). In all these graphs, the response arising from the *GLT1* locus (blue) is compared to the one monitored at the native *STL1* locus (green). Two different transformants were measured in order to verify that undesired Cas9 activity has not disrupted the HOG response.

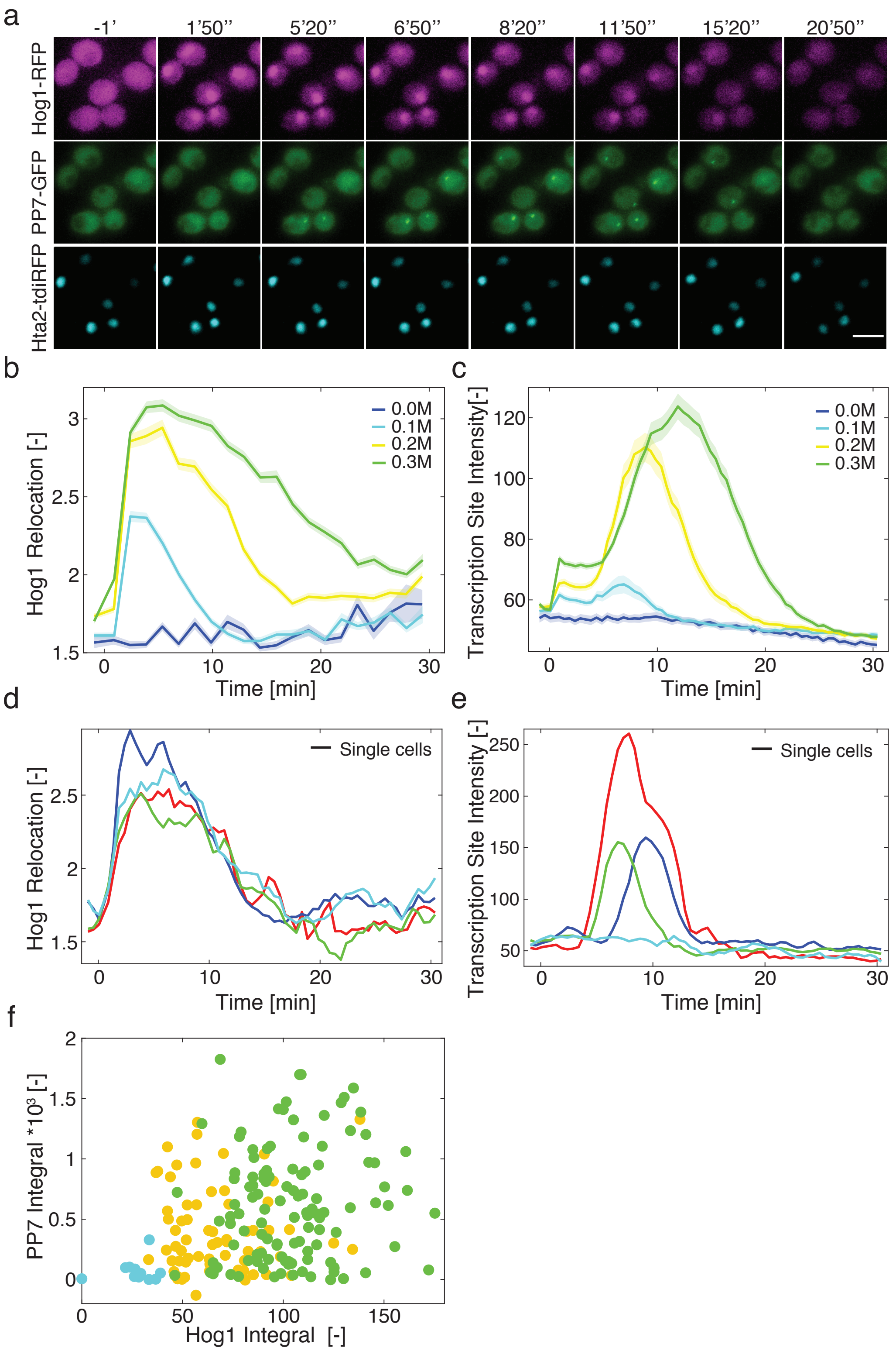


**Supplementary Figure 5. Correlating Hog1 activity and downstream transcription in the same cell.**

**a.** Thumbnails images of the strain combining a Hta2-tdiRFP nuclear marker, Hog1-mCherry and the p*STL1*-PP7 reporter. Cells were stressed with 0.2M NaCl at time 0. Scale bar is 5 $\mu$m **b. - c.** Mean dynamics of Hog1 nuclear enrichment (b) and PP7 transcription site fluorescence intensity (c) following different osmotic stresses. More than 140 cells are quantified for the inducing conditions and only 65 in the SD-full experiment. **d. - e.** Examples of single-cell traces that display similar Hog1 relocation dynamics (d) and different transcriptional responses as quantified by the p*STL1*-PP7 transcription site intensity (e). **f.** Correlation between Hog1 relocation and the PP7 output measured by their integrals. Each dot corresponds to a single-cell measurement.

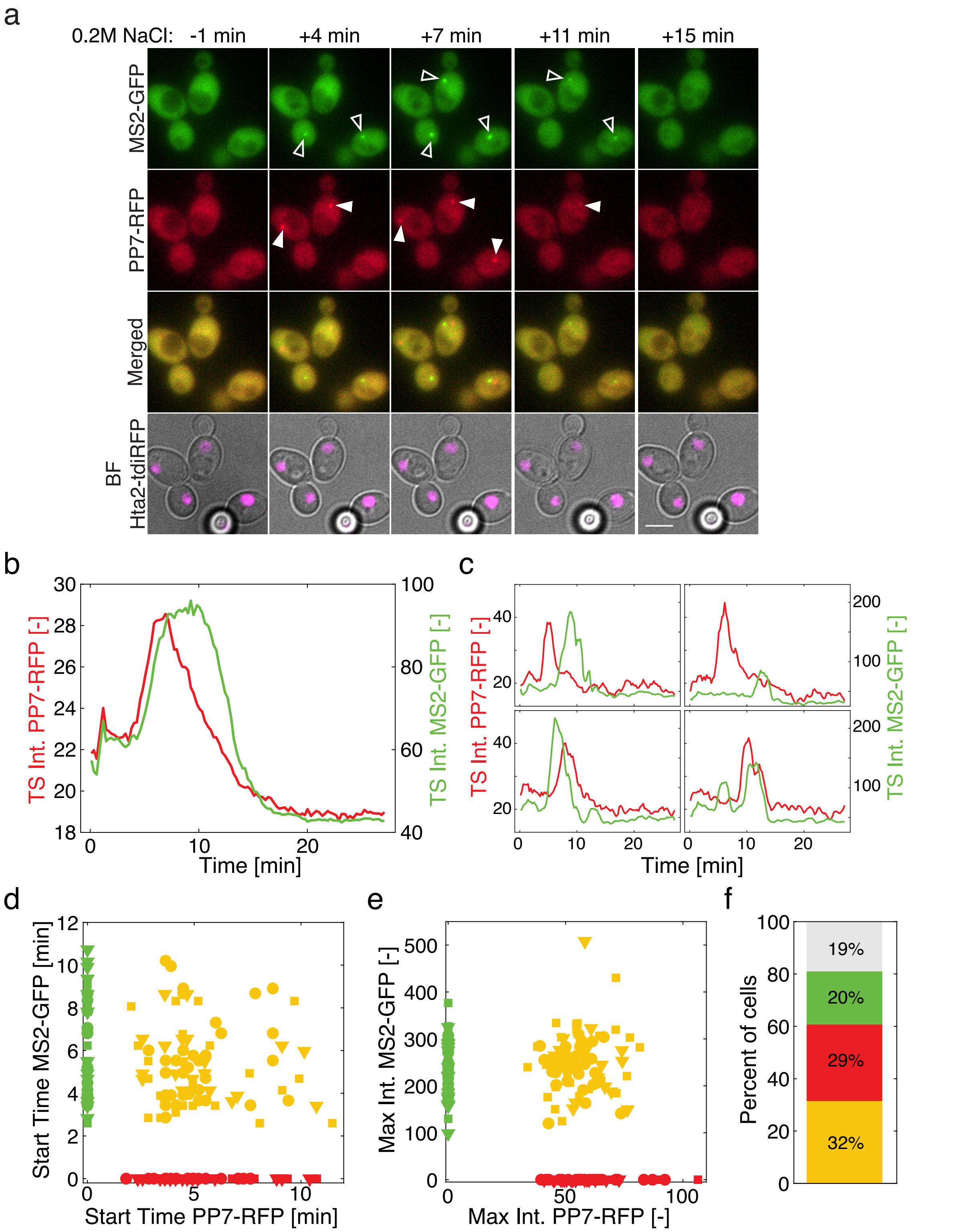


**Supplementary Figure 6. Monitoring p*STL1*-induced transcription from two identical loci in diploids.**

**a.** Thumbnails of diploid cells bearing the MS2-GFP and PP7-mCherry reporter systems monitoring the induction of two p*STL1* in the same cell following 0.2M NaCl stress. Open arrowheads (MS2sl) and closed arrowheads (PP7sl) highlight the stochastic activation of the transcription within a cell. Scale bar 5$\mu$m. **b.** Transcription site intensity from the p*STL1* promoter monitored with the MS2 (green) or the PP7 (red) systems. The low signal provided by the PP7-mCherry assay and bleaching of this FP can explain the discrepancy between the two reporter systems. **c.** Examples of single cell traces where the activation of both *STL1* promoters was detected. **d.** and **e.** Scatter plots representing the Start Times (d) and the Maximum Intensity (e) of the PP7 and MS2 assays. The data from three different experiments (rounds, squares and triangles) are combined. Cells where only the MS2 system activation was detected are plotted in green, while cells with only PP7 TS are in red. Cells where both systems where detected are in yellow. **f.** Mean percentage of cells over the three experiments where both promoters (yellow) or only one promoter (green / MS2 and red / PP7) or no transcription (gray) was detected.

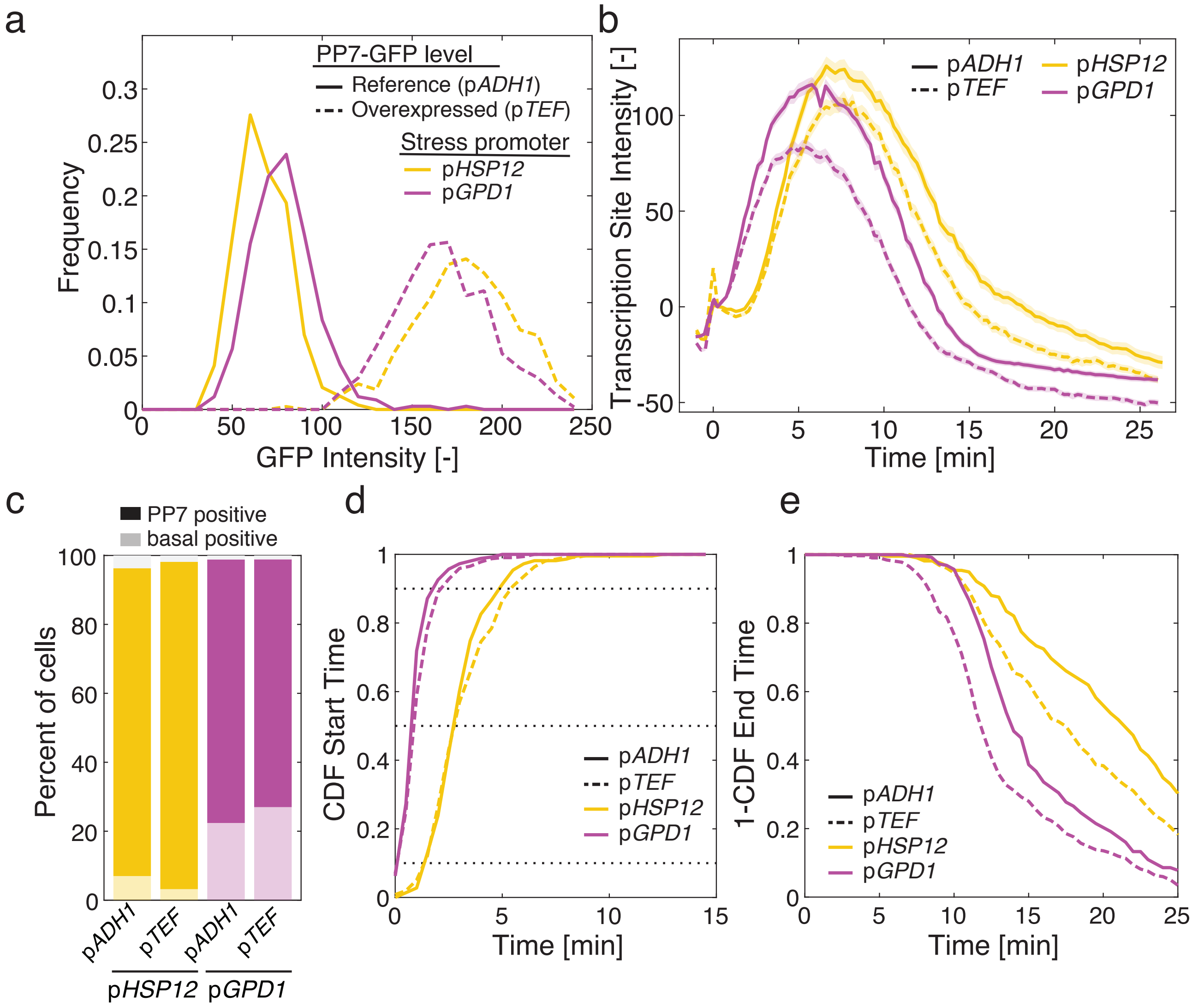


**Supplementary Figure 7. Testing the effect of the overexpression of PP7-GFP on the transcription site measurements.**

**a.** Comparison of the initial GFP fluorescence for the PP7-GFPenvy expressed from the p*ADH1* (reference) or p*TEF* promoter (overexpression) which leads to a 3-fold higher fluorescence. **b.** Mean transcription site intensity following a 0.2M NaCl stress. **c.** Percentages of PP7 positive cells. Cells displaying a TS before time point 4 (basal positive) are displayed in a lighter color. **d.** Cumulative distribution of Start Times. **e.** One minus the cumulative distribution function of End Times. The lower expression level of the PP7-GFP by the reference p*ADH1* promoter improves the detection efficiency of the TS.

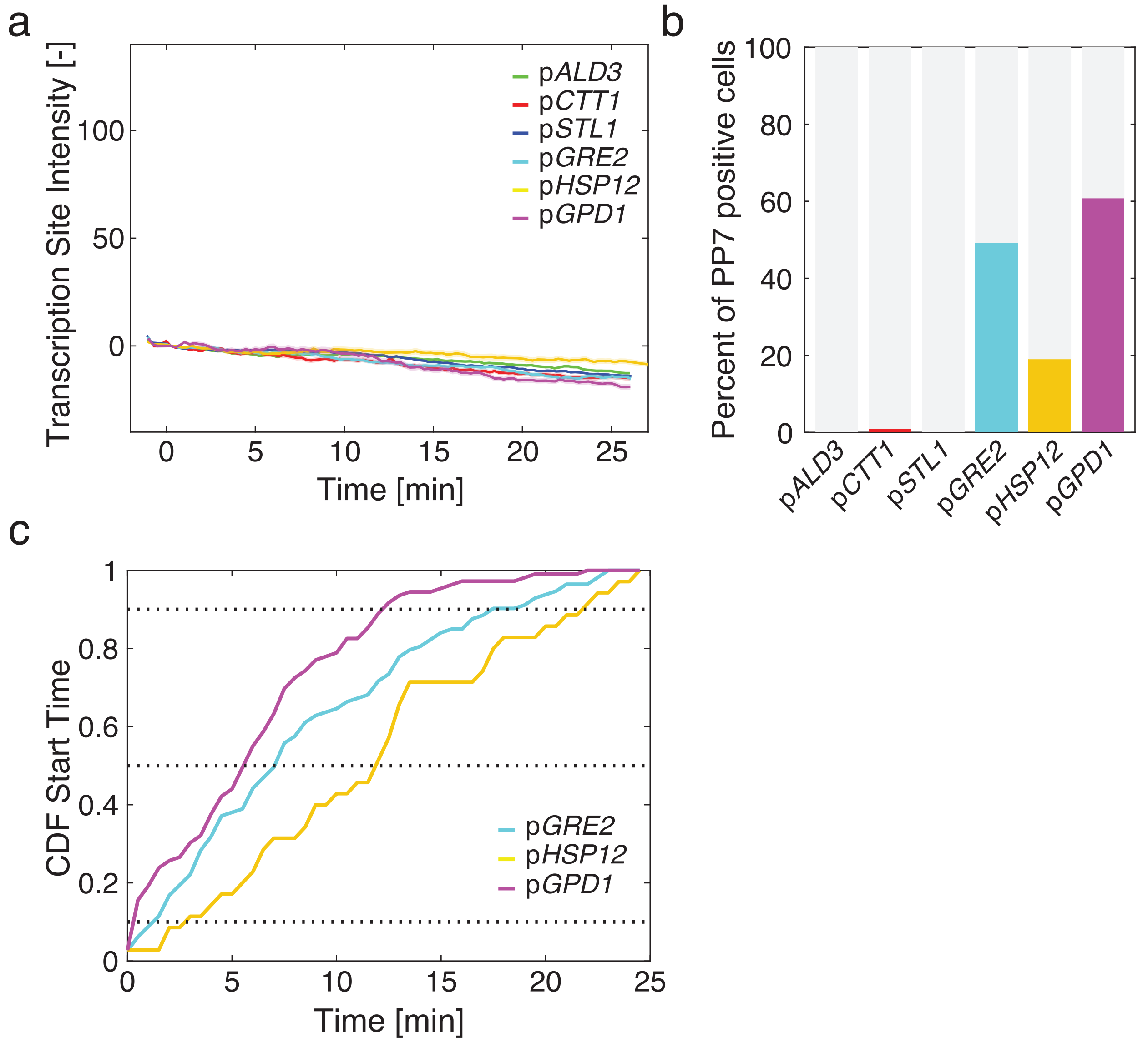


**Supplementary Figure 8. HOG promoters with basal expression level.**
**a.** Average transcription site intensity following an SD-full addition. The solid line represents the mean ratio and the shaded area represents the s.e.m. of at least 120 cells.
**b.** Percentages of PP7 positive cells during the entire SD-full time-lapse experiment in a.
**c.** Cumulative distribution function of Start Times for the promoters displaying more than 10% of expressing cells in the SD-full time-lapse experiment.

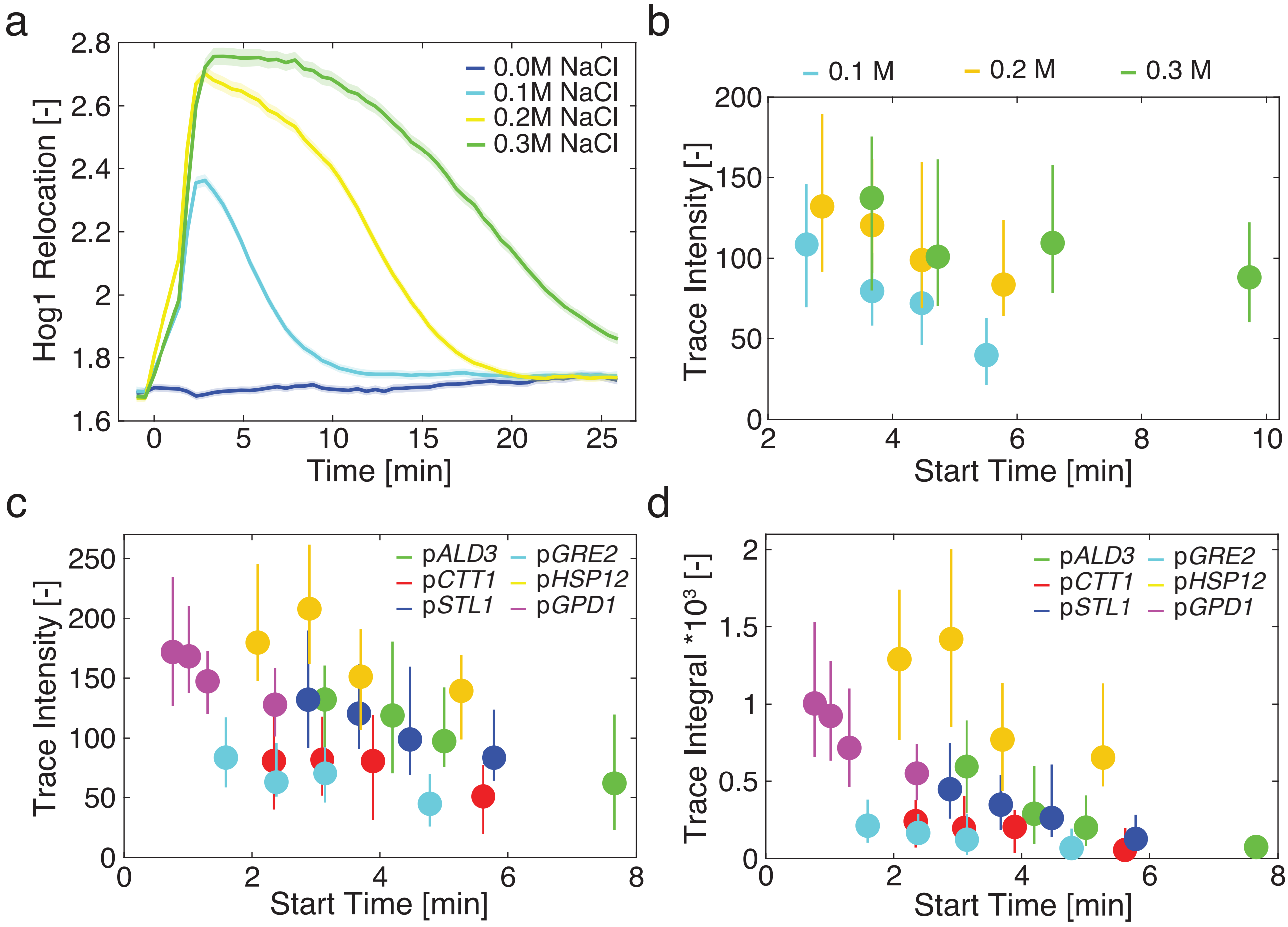


**Supplementary Figure 9. Negative correlation between Start Time and transcriptional output for all HOG promoters.**
**a.** Dynamics of Hog1 nuclear enrichment following hyper-osmotic stress. The mean ratio of nuclear over cytoplasmic fluorescence of Hog1-GFP for more than 250 cells is plotted as function of time. The shaded area represents the s.e.m. **b.** The population of p*STL1*-PP7 positive cells is split in four quartiles based on their Start Time. The median (●) and $25^{th}$ to $75^{th}$ percentiles (line) of the intensity of the PP7 trace is plotted for each quartile. **c.- d.** Plot of the Start Time versus the Trace intensity (c) or Trace integral (d) for all the PP7 reporter strains following a 0.2M NaCl stress. The population of responding cells is split in four quartiles based on their Start Time. The median (●) and $25^{th}$ to $75^{th}$ percentiles (line) of the Trace Intensity (c) or Trace Integral (d) are plotted for each quartile.

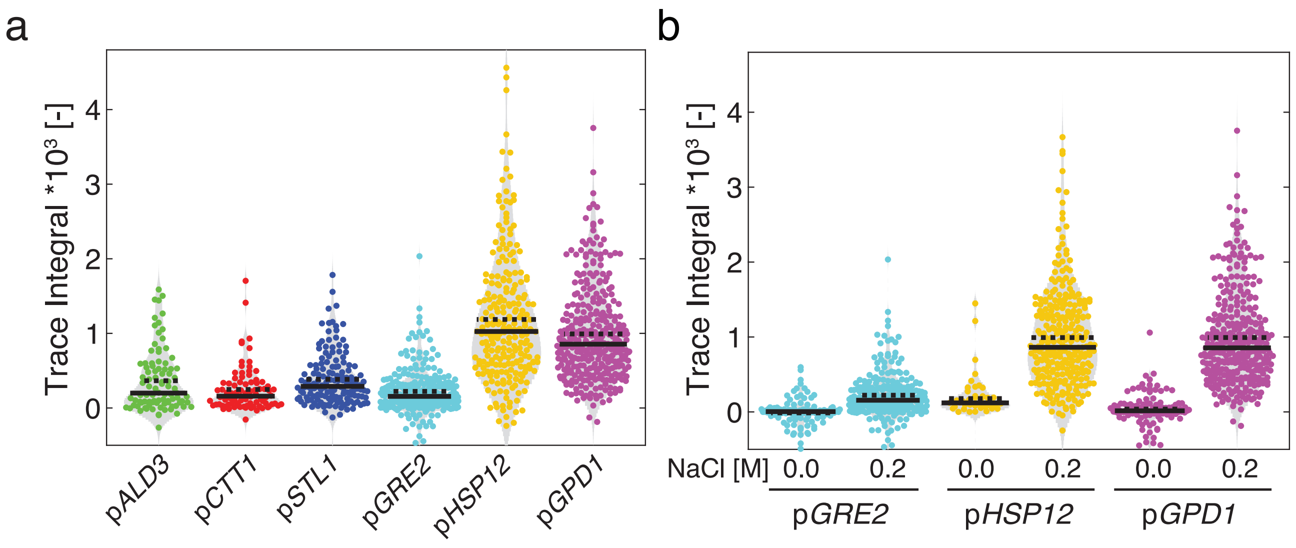


**Supplementary Figure 10. Trace integral of HOG promoters.**
**a.** Violin plots of the Trace integral of osmostress promoters response after 0.2M NaCl treatment. **b.** Violin plots of the Trace integral of basal level positive osmostress promoters response after SD-full (0.0M) or 0.2M NaCl treatment. For both graphs, each dot represents the data of a single cell, the full line the median of the response and the dashed line the mean of the response.

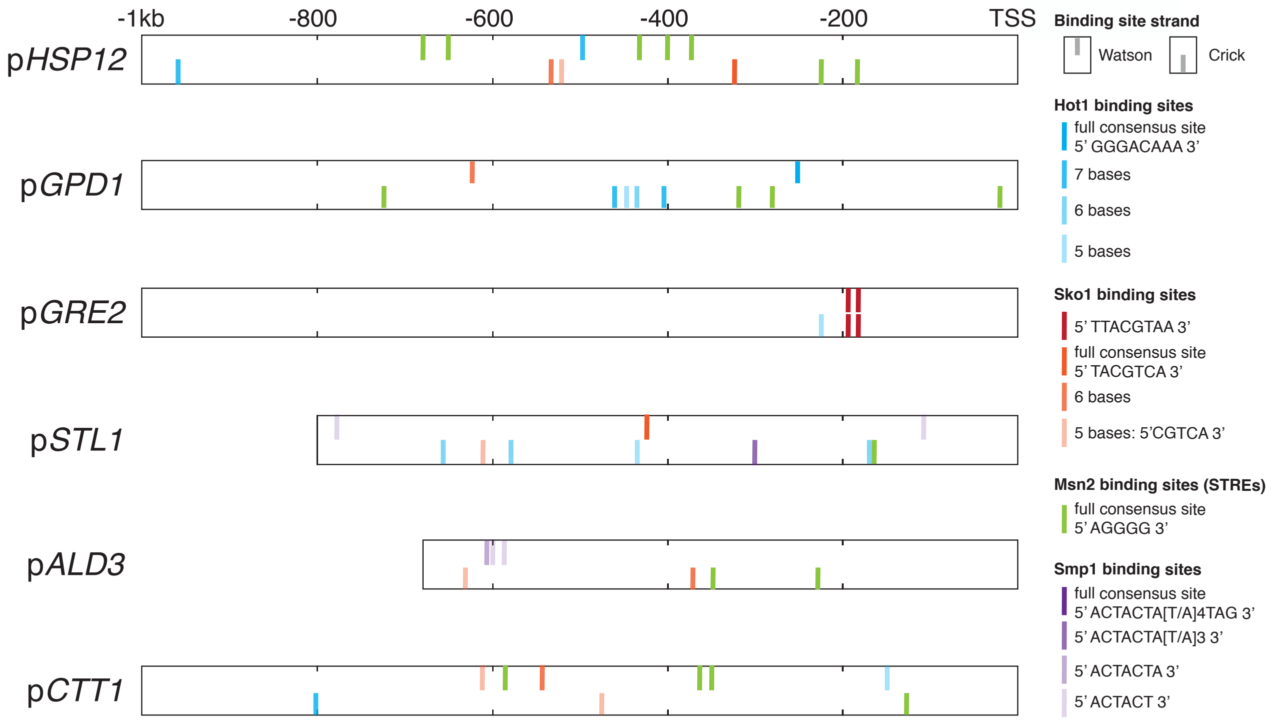


**Supplementary Figure 11. Stress promoter architecture.**
Consensus binding sites of Hot1[6], Sko1[7], Msn2/4[8], and Smp1[9] and some deviations from these consensus sequences have been mapped on the six promoters used in this study.

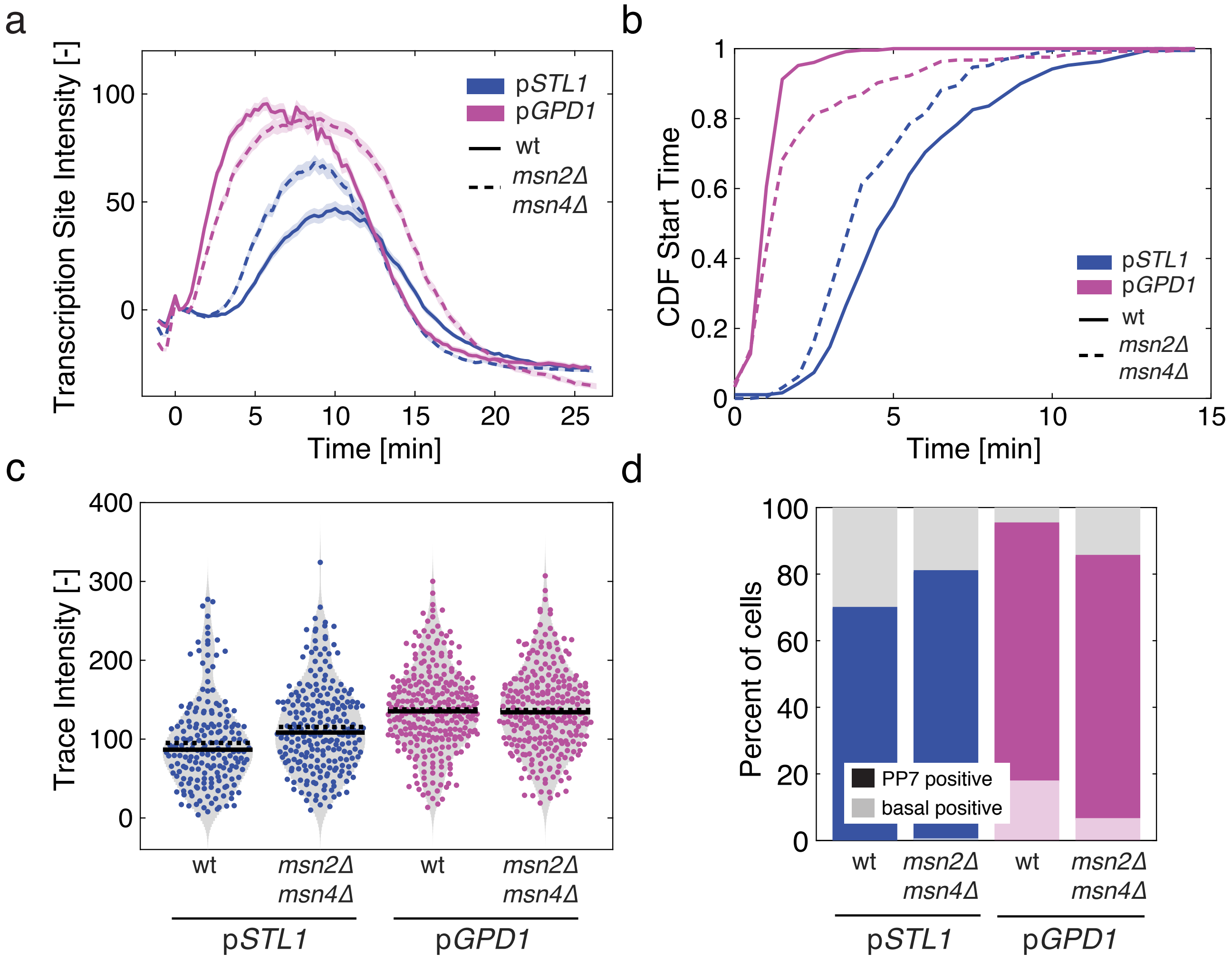


**Supplementary Figure 12. Analysis of transcription dynamics in the *msn2Δmsn4Δ* mutant**

**a.** Transcription site intensity of WT (solid line) and *msn2Δmsn4Δ* (dashed line) bearing the p*STL1*-PP7 (blue) or the p*GPD1*-PP7 (magenta) reporters following a 0.2M NaCl stress. **b.** Cumulative distribution of Start Time for the two promoters in the WT and mutant backgrounds for cells that induce transcription after time zero. **c.** Violin plots of the trace intensity (maximum of the TS during the transcription period) after stimulation by 0.2M NaCl. Each dot represents the value calculated from a single cell. The solid line is the median and the dashed line the mean of the population. **d.** Percentages of cells where a PP7 TS site was detected. The light shaded area represents the percentage of PP7 positive cells before the stimulus was added (basal transcription).

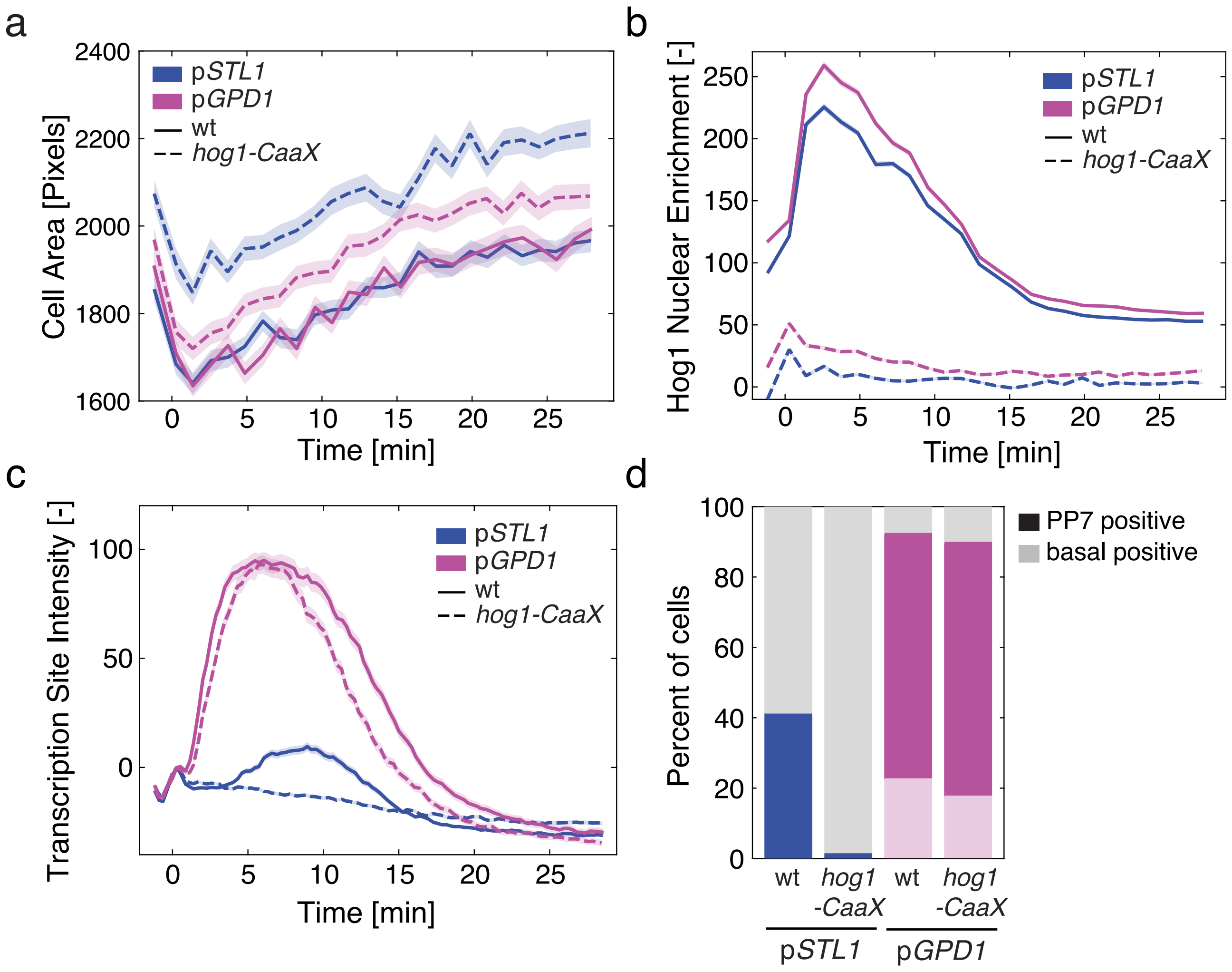


**Supplementary Figure 13. Impact of Hog1 anchoring at the plasma membrane of p*STL1* and p*GPD1* activation.**
**a. - c.** Dynamics of cell size adaptation (a), Hog1-mCherry nuclear enrichment (b) and Transcription Site intensity (c) in cells expressing either freely diffusing (wt, solid line) or membrane anchored Hog1 via a CaaX motif (dashed line) for the two transcriptional reporters p*STL1*-PP7 (blue) and p*GPD1*-PP7 (magenta) upon 0.2M NaCl stress. **d.** Percentage of cells where a PP7 TS site was detected. The light shaded area represents the percentage of PP7 positive cells before the stimulus was added (basal transcription).

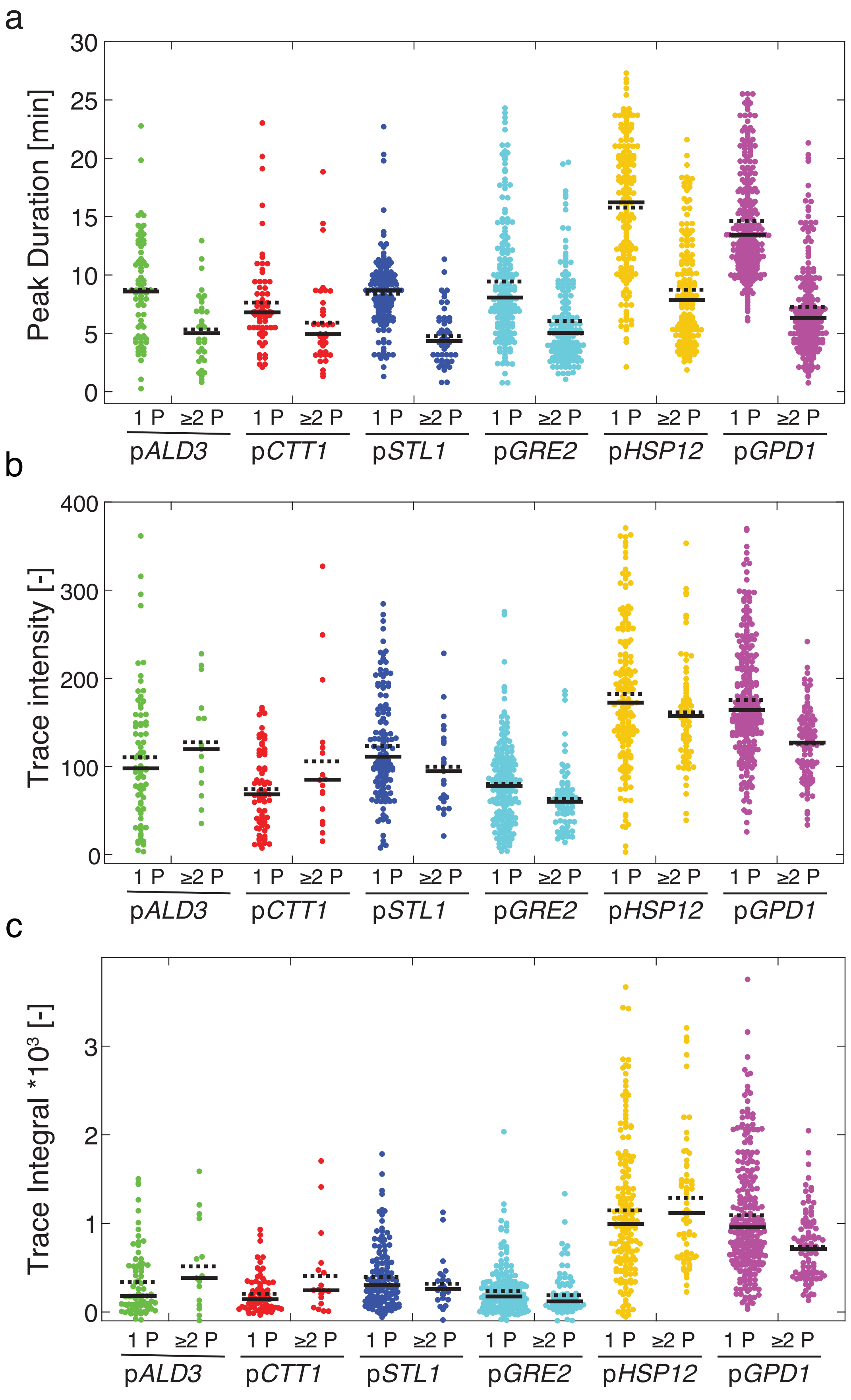


**Supplementary Figure 14. Peak analysis of osmostress-promoter in response to 0.2M NaCl.**
**a. - c.** Violin plots of the Peak Duration (a), Trace Intensity (b) and Trace Integral (c) for the different PP7 reporter strains stresses with 0.2 M NaCl. The population of cells was split between cells displaying one peak and cells where two peaks or more were detected. Each dot represents the value calculated for a single peak (a) or single cell (b and c). The solid line is the median and the dashed line the mean of the population.

**Supplementary Movies Legends**

**Supplementary Movie 1: Time lapse movie of the p*STL1*-PP7 reporter strain.**
The left image is a maximum intensity projection of a Z-stack in the green channel allowing to image the PP7-GFP and visualizing the presence of transcription sites as bright foci. The central image is the red channel image, representing the fluorescence of the Hta2-mCherry nuclear tag. The right image is a merged image between the green and red channels. The number in the upper right corner indicates the time in minutes. Cells are stressed with 0.2M NaCl at time 0.

**Supplementary Movie 2: Time lapse movie of a diploid p*STL1*-PP7 / p*STL1*-MS2 reporter strain.**
The leftmost image is a maximum intensity projection of a Z-stack in the green channel allowing to image the MS2-GFP and visualizing the presence of transcription sites as bright foci. The center-left image is a maximum intensity projection of a Z-stack in the red channel allowing to image the PP7-mCherry and visualizing the presence of transcription sites as bright foci. The center-right image is the far-red channel image representing the fluorescence of the Hta2-tdiRFP nuclear tag. The rightmost image is a merged image between the green and red channels (PP7 and MS2) allowing to observe to which extent the induction of two pSTL1 correlate in the same cell. The number in the lower right corner indicates the time in minutes. Cells are stressed with 0.2M NaCl at time 0.

**Supplementary Movie 3: Time lapse movie of the p*GPD1*-PP7 reporter strain.**
The left image is a maximum intensity projection of a Z-stack in the green channel allowing to image the PP7-GFP and visualizing the presence of transcription sites as bright foci. Note the presence of some transcription site in absence of stimulus in the first frames of the movie. The central image is the red channel image, representing the fluorescence of the Hta2-mCherry nuclear tag. The right image is a merged image between the green and red channels. The number in the upper right corner indicates the time in minutes. Cells are stressed with 0.2M NaCl at time 0.

**Supplementary Movie 4: Time lapse movie of the p*STL1*-PP7 in a step experiment.**
The leftmost image is a maximum intensity projection of a Z-stack in the green channel allowing to image the PP7-GFP and visualizing the presence of transcription sites as bright foci. The background fluorescence in the image allows to follow NaCl concentration changes in the flow channel. Higher fluorescence is indicative of lower NaCl concentrations. The center-left image is the red channel image and allows to follow the changes in Hog1 nuclear localization. The center-right image is the far-red channel image representing the fluorescence of the Hta2-tdiRFP nuclear tag. The rightmost image is a merged image between the green and red channels (PP7 and Hog1). The number in the lower right corner indicates the time in minutes from the start of the experiment.

**Supplementary Movie 5: Time lapse movie of the p*STL1*-PP7 in a pulse experiment.**
The leftmost image is a maximum intensity projection of a Z-stack in the green channel allowing to image the PP7-GFP and visualizing the presence of transcription sites as bright foci. The background fluorescence in the image allows to follow NaCl concentration changes in the flow channel. Higher fluorescence is indicative of lower NaCl concentrations. The center-left image is the red channel image and allows to follow the changes in Hog1 nuclear localization. The center-right image is the far-red channel image representing the fluorescence of the Hta2-tdiRFP nuclear tag. The rightmost image is a merged image between the green and red channels (PP7 and Hog1). The number in the upper right corner indicates the time in minutes from the start of the experiment.

**Supplementary Movie 6: Time lapse movie of the p*STL1*-PP7 in a ramp experiment.**
The leftmost image is a maximum intensity projection of a Z-stack in the green channel allowing to image the PP7-GFP and visualizing the presence of transcription sites as bright foci. The background fluorescence in the image allows to follow NaCl concentration changes in the flow channel. Higher fluorescence is indicative of lower NaCl concentrations. The center-left image is the red channel image and allows to follow the changes in Hog1 nuclear localization. The center-right image is the far-red channel image representing the fluorescence of the Hta2-tdiRFP nuclear tag. The rightmost image is a merged image between the green and red channels (PP7 and Hog1). The number in the lower right corner indicates the time in minutes from the start of the experiment.
